# The rise of diversity in metabolic platforms across the Candidate Phyla Radiation

**DOI:** 10.1101/2019.12.18.881540

**Authors:** Alexander L. Jaffe, Cindy J. Castelle, Paula B. Matheus Carnevali, Simonetta Gribaldo, Jillian F. Banfield

## Abstract

A unifying feature of the bacterial Candidate Phyla Radiation (CPR) is a limited and highly variable repertoire of biosynthetic capabilities. However, the distribution of metabolic traits across the CPR and the evolutionary processes underlying them are incompletely resolved. Here, we selected ∼1,000 genomes of CPR bacteria from diverse environments to construct a robust internal phylogeny that was consistent across two unlinked marker sets. Mapping of glycolysis, the pentose phosphate pathway, and pyruvate metabolism onto the tree showed that some components of these pathways are sparsely distributed and that similarity between metabolic platforms is only partially predicted by phylogenetic relationships. To evaluate the extent to which gene loss and lateral gene transfer have shaped trait distribution, we analyzed the patchiness of gene presence in a phylogenetic context, examined the phylogenetic depth of clades with shared traits, and compared the reference tree topology with those of specific metabolic proteins. While the central glycolytic pathway in CPR is widely conserved and has likely been shaped primarily by vertical transmission, there is evidence for both gene loss and transfer especially in steps that convert glucose into fructose 1,6-bisphosphate and glycerate 3P into pyruvate. Additionally, the distribution of Group 3 and Group 4-related NiFe hydrogenases is patchy and suggests multiple events of ancient gene transfer. Overall, patterns of gene gain and loss, including acquisition of accessory traits in independent transfer events, may have been driven by shifts in host-derived resources and led to sparse but varied genetic inventories.

## INTRODUCTION

Metagenomics approaches have been extremely fruitful in the discovery of new lineages across the tree of life (Anantharaman et al., 2016; Brown et al., 2015; Parks et al., 2017; Rinke et al., 2013). Through increased sampling of diverse environments, genomes recovered from poorly represented or novel groups have elucidated evolutionary processes contributing to bacterial and archaeal diversity (Adam et al., 2017; Castelle et al., 2018; Spang et al., 2015). These studies have resolved metabolic capacities in little known or unknown organisms and documented evidence for processes like lateral gene transfer that shape the distribution of metabolic capacities over various lineages.

The Candidate Phyla Radiation, a large group of bacterial lineages lacking pure isolate cultures, are one group primarily defined through genome-resolved metagenomics (Brown et al., 2015; Luef et al., 2015). While estimates vary depending on the methods used (Hug et al., 2016; Parks et al., 2018), CPR are predicted to constitute a significant portion of bacterial diversity that is distinct and divergent from other bacterial groups (Zhu et al., 2019). Additionally, CPR bacteria generally have relatively small genome and cell sizes, extremely reduced genomic repertoires, and often lack the capacity to synthesize lipids (Brown et al., 2015; Kantor et al., 2013; Luef et al., 2015). The CPR may have diverged early from other bacteria and subsequently diversified over long periods of time, or they may have arisen via rapid evolution involving genome streamlining/reduction (Castelle and Banfield, 2018). Arguing against recent diversification from other bacteria are the observations that CPR do not share genomic features associated with recent genome reduction, have uniformly small genomes, cluster independently from other metabolically reduced symbionts, and possess metabolic platforms consistent with projections for the anaerobic environment of the early Earth (Castelle and Banfield, 2018; Méheust et al., 2019; Schönheit et al., 2016).

Recently, an analysis of entire proteomes showed that genetic capacities encoded by CPR are combined in an enormous number of different ways, yet those combinations tend to recapitulate inferred phylogenetic relationships between groups (Meheust et al. 2019). These analyses also revealed that some lineages have relatively minimal core gene sets compared to other CPR (Castelle et al., 2018; Méheust et al., 2019), suggesting variation in the degree of genome reduction across the radiation. Additionally, previous work has shown that lateral gene transfer probably underlies distributions of specific protein families in the CPR, including RuBisCO (Jaffe et al., 2016, 2019). The observation that CPR also encode genes for nitrogen, hydrogen, and sulfur compound transformations at low frequency (Castelle et al., 2018; Danczak et al., 2017; Wrighton et al., 2012, 2014, 2016) raises the possibility that these capacities may also have been shaped by lateral transfer. Overall, the extent to which lateral transfer, genomic loss, and vertical transfer have interacted to shape evolution of metabolic repertoires in the CPR is still unknown (Castelle and Banfield, 2018).

Here, we integrate insights from CPR genomes from diverse environments with a robustly-resolved internal phylogeny to investigate the processes governing the evolution of metabolic pathways in CPR bacteria. A key aspect of our approach was the development of custom cutoffs for HMM-based metabolic annotation that are sensitive to the divergent nature of CPR proteins. We investigated central carbon metabolism (glycolysis and the pentose phosphate pathway), hypothesizing that these pathways may be primarily shaped by vertical inheritance, as well as sparsely distributed traits (nitrogen, hydrogen, sulfur metabolism) that we predicted were shaped by lateral transfer. Mapping of metabolic capacities onto the reconstructed tree and gene-species tree reconciliations showed that a mixture of vertical inheritance, gene loss, and lateral transfer have differentially shaped the distribution of functionally linked gene sets. Information about the evolution of gene content may help to shed light on evolutionary scenarios that broadly shaped the characteristics of extant CPR bacteria.

## RESULTS

### Concatenated ribosomal protein and RNA polymerase subunits outline a robust internal phylogeny for CPR

We gathered approximately 2,300 curated CPR genomes from diverse environments, including both previously published and newly assembled sequences (Materials and Methods). Quality filtration of this curated genome set at ≥70% completeness and ≤10% contamination and subsequent de-replication yielded a non-redundant set of 991 CPR genomes for downstream phylogenetic and metabolic analysis (Supp. Table 1). To improve recovery of phylogenetic markers from the collected set of genomes, we combined visualization of HMM bitscores with a phylogenetic approach to set sensitive, custom thresholds for two independent sets of markers composed of 16 syntenic ribosomal proteins (rp16) and the two RNA polymerase subunits (RNAp) (Materials and Methods). Phylogenies based on these two marker sets were generally congruent for deep relationships within the CPR, with both trees supporting the monophyly of the Microgenomates and Parcubacteria superphyla. Mapping of presence/absence of ribosomal protein L9 (rpL9) onto the tree confirmed that the Microgenomates, along with the Dojkabacteria and Katanobacteria, are part of a larger clade lacking this protein, as suggested previously (Brown et al., 2015). Similarly, mapping of the ribosomal protein L1 (rpL1) onto the tree was consistent with the finding that a monophyletic group of Parcubacteria lack this enzyme (Brown et al., 2015). Our results also suggested the presence of four generally well-supported (≥95% ultrafast bootstrap in three of four cases), monophyletic subgroups within the Parcubacteria (Fig. 1a, Parcubacteria 1-4). Although internal relationships between these subgroups varied slightly between trees (Supp. Fig. 3), in both cases Parcubacteria 1 (comprised of 9 lineages) was relatively deeply-rooting within the superphylum, whereas Parcubacteria 4 (10 lineages) was the most shallow rooting (Fig. 1a). Using a diverse bacterial outgroup of around ∼170 genomes, we also show that Dojkabacteria (WS6), Katanobacteria (WWE3), Peregrinibacteria, Kazanbacteria, and Berkelbacteria are among the most deeply-rooting clades outside the established superphyla, consistent with previous tree reconstructions made from smaller genome sets (Anantharaman et al., 2016; Hug et al., 2016) (Fig. 1a).

**Figure 1.**
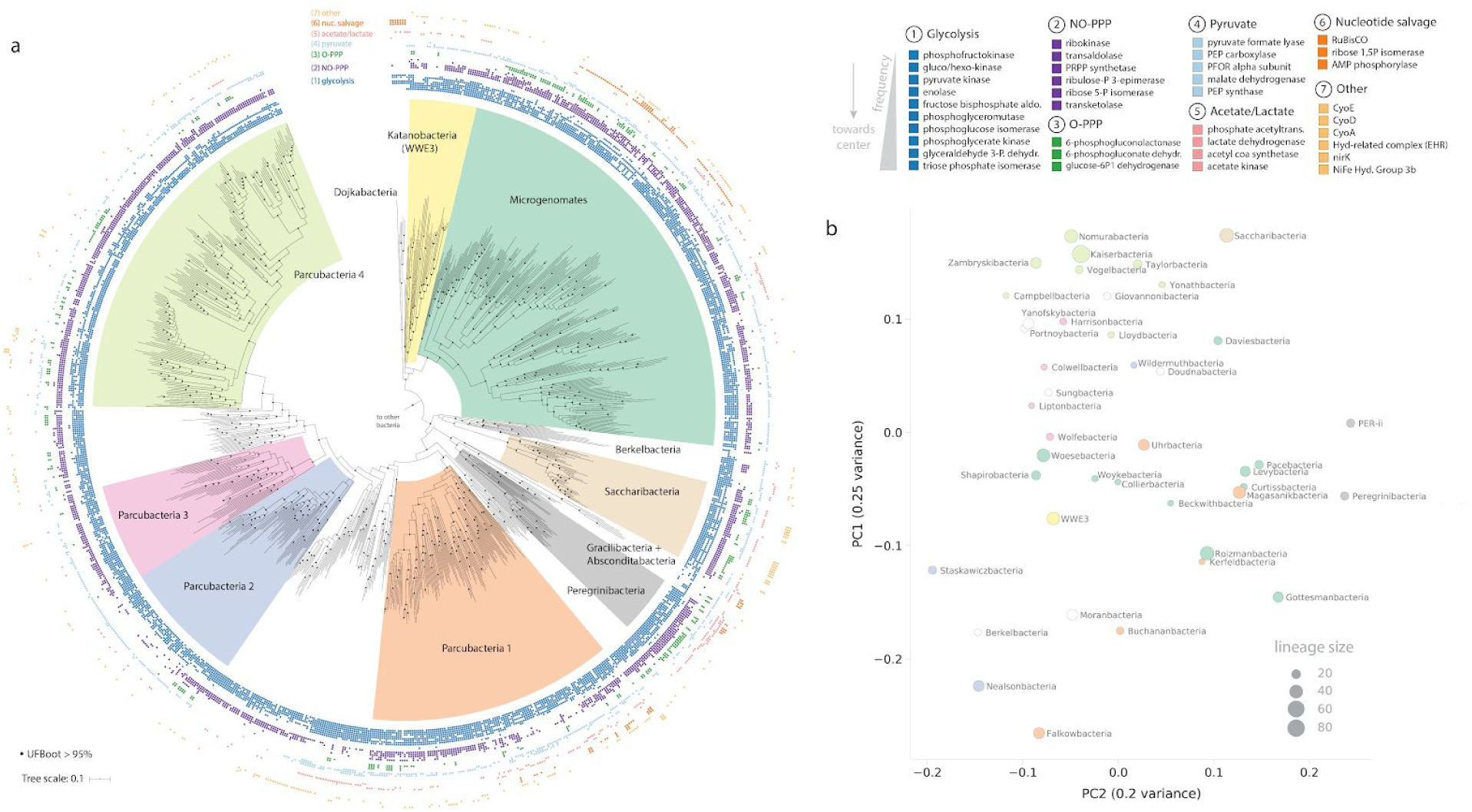
Phylogenetic relationships and metabolic similarity among Candidate Phyla Radiation bacteria. **a)** Maximum-likelihood tree based on the concatenated set of 16 ribosomal proteins (1427 amino acids, LG+R10 model). Scale bar represents the average number of substitutions per site. Monophyletic subgroups within the Parcubacteria also supported in the concatenated RNA polymerase tree are indicated as Parcubacteria 1-4. Presence/absence of a subset of targeted metabolic traits are indicated as concentric rings. Abbreviations: aldo., aldolase; dehydr., dehydrogenase, PRPP, phosphoribosylpyrophosphate, PEP, phosphoenolpyruvate; PFOR, pyruvate:ferredoxin oxidoreductase; acetyltrans., acetyltransferase; Hyd, hydrogenase. Fully annotated trees with all included lineages are available in Supp. Fig. 3. **b)** Principal coordinates analysis describing similarity between metabolic platforms of CPR lineages with 8 or more representative genomes.

### CPR bacteria encode variable and overlapping metabolic repertoires

A major objective of this study was to leverage the constructed reference tree of the CPR to evaluate the distribution and combinations of capacities across the radiation. While CPR bacteria lack some core biosynthetic capacities, they do in fact possess numerous metabolic capacities involved in carbon, hydrogen, and possibly sulfur and nitrogen cycling (Castelle et al., 2018; Danczak et al., 2017; Kantor et al., 2013; Wrighton et al., 2012). We focused on these traits for our subsequent analysis, reasoning that they are most likely to impact the ability of CPR bacteria to derive energy from organic compounds and contribute to biogeochemical transformations in conjunction with their hosts and other community members. To overcome the challenges inherent to metabolic annotation of divergent lineages and minimize the chance of false negatives, and we extended our custom HMM thresholding approach to the selected set of traits (Materials and Methods, Supp. Figure 2) and mapped the resulting binary presence/absence profiles for specific functionalities onto the reconstructed rp16 tree (Figure 1). Our results reveal a striking patchiness in distribution of traits across the CPR radiation, even for some components of glycolysis and the pentose phosphate pathway (Fig. 1a). Intriguingly, traits involved in pyruvate, acetate, oxygen, nucleotide, and hydrogen/sulfur metabolism exhibited sparse but also wide distributions across distantly related groups, consistent with prior analysis of individual genomes (Castelle et al., 2018; Wrighton et al., 2012) (Fig. 1a). Looking across the selected traits, there is clear variation in the overall repertoires of lineages within CPR, including some with extremely minimal metabolic complements like Dojkabacteria and Gracilibacteria. This is consistent with both observations from genomic studies of these lineages (Kantor et al., 2013) as well as more recent insights examining entire proteomes (Méheust et al., 2019).

**Figure 2.**
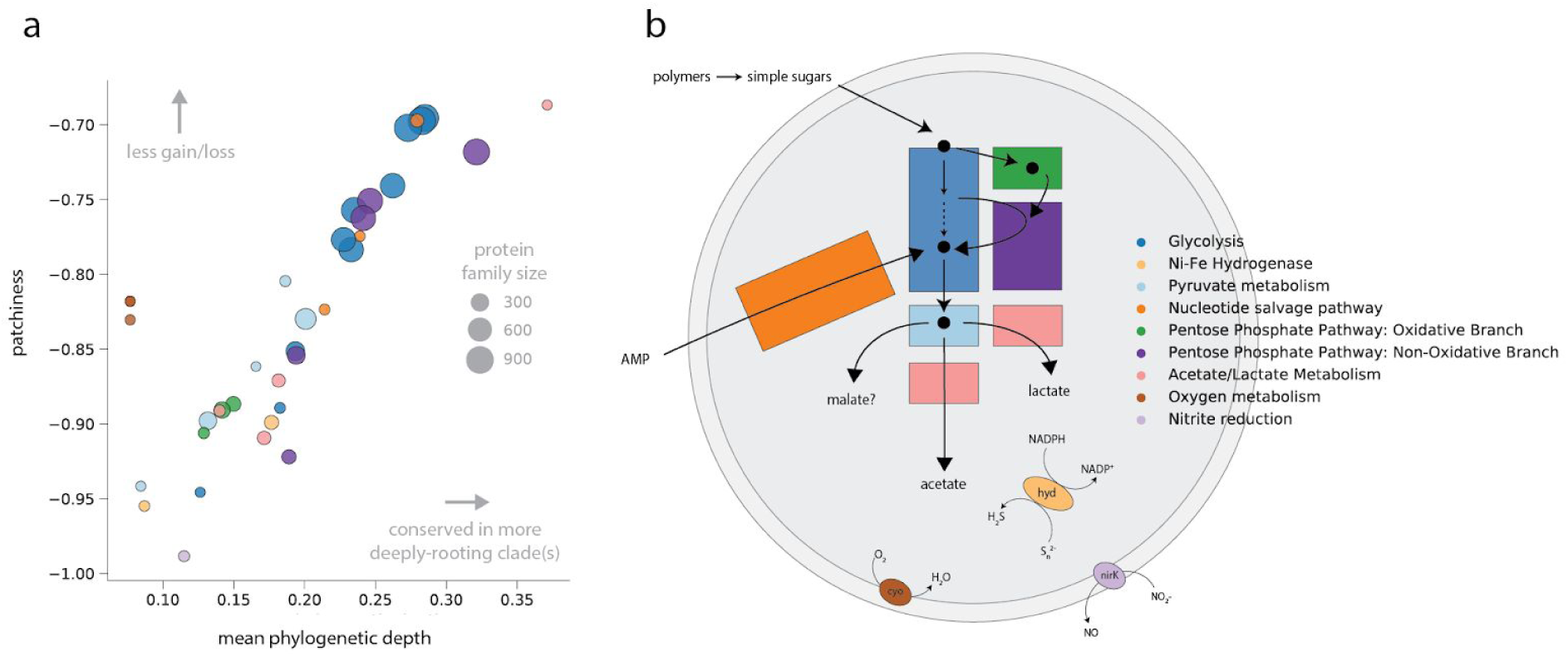
Metabolic traits encoded by CPR exhibit varying evolutionary profiles, including those in the same pathway (e.g. glycolysis genes). **a)** Evolutionary profiles generated from phylogenetic depth and patchiness of gene distributions over the rp16 topology. Each point represents a metabolic gene shaded to match the functional category/pathway in **b)**, schematic representing a generalized metabolic platform for CPR bacteria.

**Figure 3.**
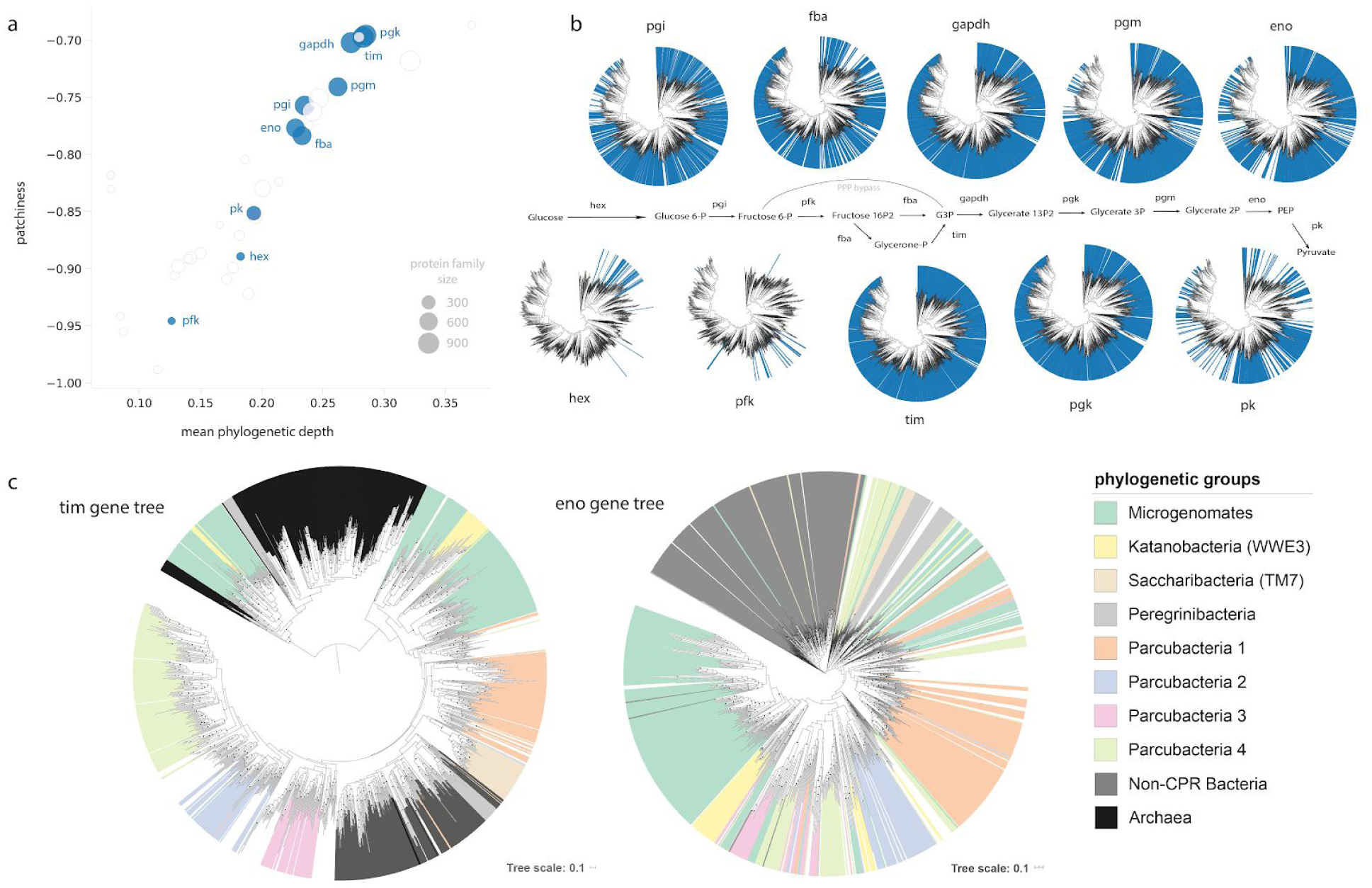
Patterns of distribution and gene trees for glycolytic enzymes in the CPR. **a)** Evolutionary profiles based on patchiness and phylogenetic depth and **b)** presence/absence profiles over the rp16 tree. **c)** Protein-specific molecular phylogenies for triose phosphate isomerase (tim) and enolase (eno). Abbreviations: hex, hexokinase; pfk, phosphofructokinase; pk, pyruvate kinase; fba, fructose bisphosphate aldolase; eno, enolase; pgi, phosphoglucose isomerase; pgm, phosphoglycerate mutase; tim, triose phosphate isomerase; gapdh, glyceraldehyde 3-phosphate dehydrogenase; pgk, phosphoglycerate kinase; G3P, glyceraldehyde 3-phosphate; PEP, phosphoenolpyruvate; PPP, pentose phosphate pathway. Scale bars represent the average number of substitutions per site.

An important open question is whether the observed combinations of biogeochemically-relevant metabolic capacities recapitulate species relationships among the CPR. To test the hypothesis that more closely related CPR lineages encode more similar combinations of metabolic capabilities, we used the distributions of the targeted traits to compute the frequency at which each trait was found within lineages. We then generated a distance matrix from the results and performed a principal coordinate analysis to visualize clustering of lineages based on the similarity of their overall metabolic platforms. We reasoned that genes missing due to genome incompleteness could impact clustering, particularly for small lineages with only several members. Thus, we restricted the analysis to those groups with at least 8 genomes. The results suggested that member lineages within some broad phylogenetic groupings were metabolically similar (e.g., Parcubacteria 3 and 4 in Figure 1b) but others clustered more closely with lineages that were distantly related. For example, lineages within the Microgenomates and Parcubacteria 1 were highly dispersed across the axes of variation, suggesting that member groups encode highly variable combinations of traits. In the future, increased availability of complete genomes will help to clarify and potentially validate these patterns.

### Functionally linked metabolic genes display different ‘evolutionary profiles’

The observation that distributions of traits are variable, sometimes patchy, and potentially decoupled from phylogenetic relatedness raises the possibility that more complex, enzyme-specific patterns might underlie processes contributing to metabolic diversity in the CPR. To address this, we drew upon trait distributions to compute two metrics - one to quantify the average branch length of clades in which a trait is conserved (phylogenetic depth) and a second, patchiness, related to the number of gains/losses of a binary trait over a tree (see Materials and Methods) (Mendler et al., 2019). These metrics were then integrated to create an ‘evolutionary profile’ for each trait. Generally, traits with a higher phylogenetic depth are expected to be conserved in more deeply-rooting clades, whereas traits with lower depth are expected to occur primarily among shallow clades. Similarly, high patchiness is expected when traits are more randomly dispersed across a given clade, whereas those with low patchiness scores are expected to be highly conserved within groups where they are present. In the CPR, we observed that phylogenetic depth generally increases with decreasing patchiness (Fig. 2a). High-depth traits also frequently corresponded to larger protein families, though several smaller protein families (phosphate acetyltransferase, AMP phosphorylase, RuBisCO) reached relatively high depths because they were conserved in deeply-rooting clades like the Dojkabacteria and Peregrinibacteria. Patchily distributed traits with low phylogenetic depth included hydrogen/sulfur metabolism, acetate/lactate metabolism, and the oxidative pentose phosphate pathway (Fig. 2). On the other hand, some traits had lower depth and were less patchily-distributed than would be predicted from the typical trend (e.g., genes involved in aerobic metabolism).

Surprisingly, metabolic genes within the same pathway often showed disparate evolutionary profiles - for example, enzymes involved in glycolysis displayed a wide range in depth and patchiness (Figure 2a). Similar patterns were observed for the nucleotide salvage pathway and non-oxidative pentose phosphate pathway. Differences in ‘evolutionary profile’ within a pathway might suggest that evolutionary histories of their component enzymes were decoupled. Previous work demonstrating that the phylogenetic tree for RuBisCO is incongruent with those of the other enzymes with which it functions supports this conclusion (Jaffe et al., 2019). We investigated two cases in more detail - first, glycolysis, as an example of a pathway with a wide range of phylogenetic depth and patchiness, and, second, NiFe hydrogenases, as an example of a function with high patchiness and low phylogenetic depth (Fig. 2a, Fig. 3a). These modules are functionally well-characterized and represent core and accessory capacities in the CPR.

### Gene trees for glycolytic enzymes reflect different patterns of gene loss and transfer

We first examined glycolysis, noticing that three enzymes from the central part of the pathway - triose phosphate isomerase (TIM), glyceraldehyde 3-phosphate (GAPDH), and phosphoglycerate kinase (PGK) were found in nearly all CPR bacteria with little to no patchiness (Fig. 2a). With the possible exception of ultra-reduced forms like the Gracilibacteria, which is represented by one complete, curated genome that completely lacks the glycolysis pathway (Sieber et al., 2019), the absence of these enzymes in a very small number of CPR genomes is likely due to missing genomic information. A second cluster, comprised of fructose bisphosphate aldolase (FBA), enolase (eno), and phosphoglucose isomerase (PGI), was generally more patchily distributed among CPR and missing in some lineages. Phosphoglycerate mutase (PGM), responsible for converting Glycerate 1,3-P2 to Glycerate 2P in lower glycolysis, fell between the two clusters - while present in deeply-rooting clades (thus, a high phylogenetic depth), it is absent in several shallow clades of Parcubacteria, possibly because these forms were too divergent to be included at the manual HMM threshold. Finally, several enzymes, including glucokinase/hexokinase, phosphofructokinase (PFK), and pyruvate kinase, exhibited profiles that were highly patchy and lower-depth among CPR lineages. Notably, these enzymes are thought to catalyze irreversible reactions and thus act as important sites of regulation for metabolic flux (Bräsen et al., 2014; Castelle et al., 2018). In particular, glucokinase/hexokinase and PFK were found very infrequently in CPR, though many CPR have the potential to bypass PFK using a metabolic shunt through the non-oxidative pentose phosphate pathway (Fig 3b) (Kantor et al., 2013). Despite the fact that some CPR encoded ROK family proteins (TIGR00744), we could not establish close phylogenetic relationships to family members functioning as putative glucokinases. Likewise, we found no evidence for the alternative versions of ADP-dependent glucokinase/phosphofructokinase employed in the modified glycolytic pathways of some archaea (PF04587) (Tuininga et al., 1999).

To further investigate specific processes impacting the evolution of glycolytic enzymes, we reconstructed single-protein molecular phylogenies and performed gene-species tree reconciliations. We reasoned that enzymes whose evolutionary histories were shaped primarily by vertical transfer paired with genomic loss, rather than transfer, would display phylogenetic patterns roughly congruent with our resolved species tree, whereas those impacted by horizontal transfer (with either CPR or non-CPR groups) would exhibit relationships incongruent with the resolved species tree. Gene trees for well-conserved glycolytic capacities like TIM and PGK generally recapitulated phylogenetic groupings at a coarse level (Fig. 3c, Supp. Fig. 5). However, even for these enzymes, inconsistencies with the species tree were present - for example, some triose phosphate isomerase (TIM) sequences from the Microgenomates, Katanobacteria, and Peregrinibacteria clustered with archaeal reference sequences (Fig. 3c). These results were replicated across multiple genomes and the phylogenetic associations of some scaffolds were verified to rule out mis-binning. Similarly, in the enolase phylogeny, large, monophyletic clusters representing sequences from the Microgenomates and Parcubacteria 1 were resolved; however, other sequences from the Microgenomates and many from Parcubacteria 3 and 4 fell into smaller, fragmented groups that clustered with more distantly related lineages. Gene trees for other glycolytic enzymes displayed a range of patterns (Supp. Fig. 4). On the whole, gene-species tree inconsistencies suggest that lateral gene transfer, either between CPR and other taxa or among different CPR, has also impacted the evolution of glycolytic enzymes alongside the gene loss apparent from presence/absence profiles (Fig. 3a).

**Figure 4:**
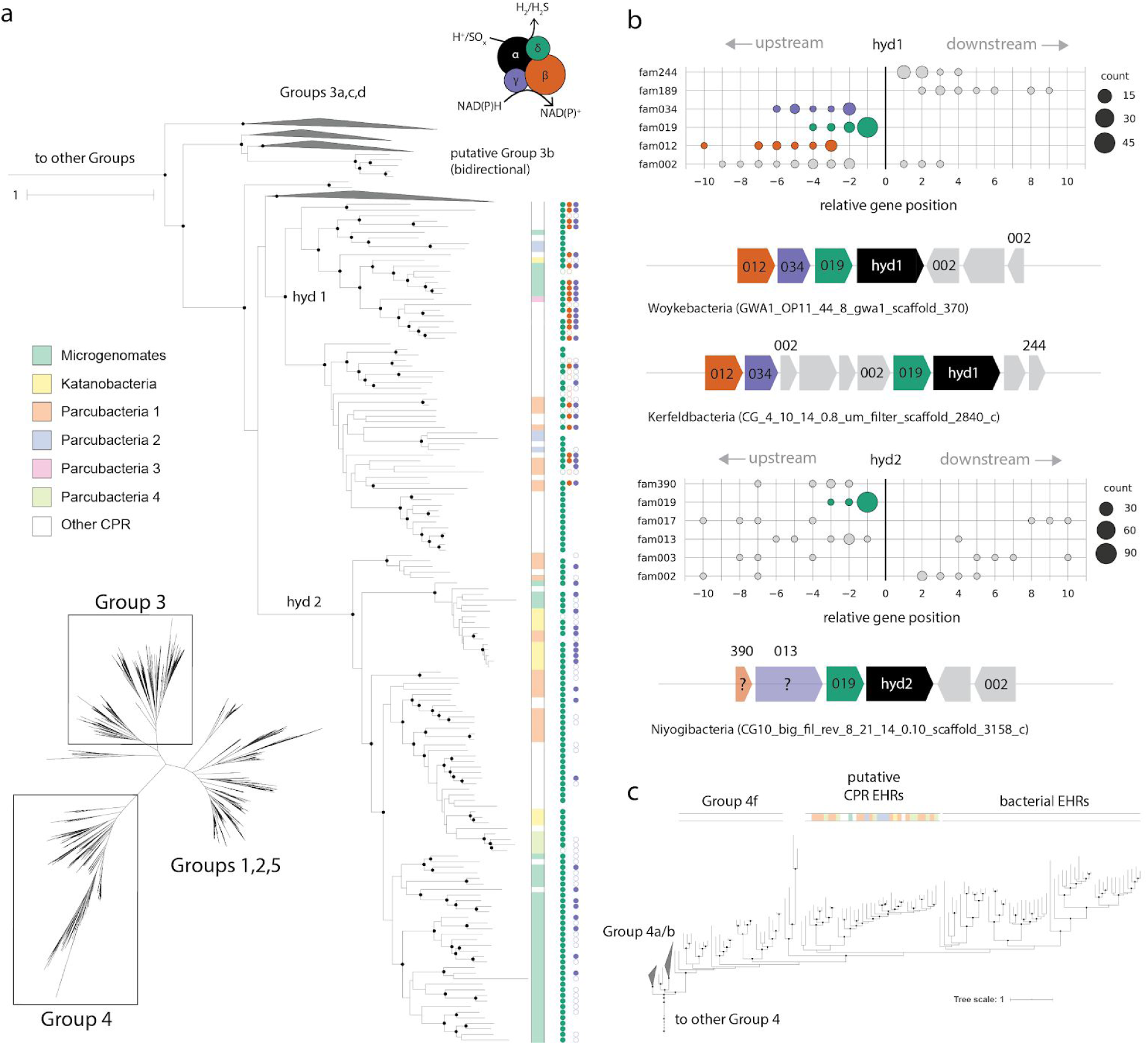
NiFe hydrogenase enzymes encoded by CPR. **a)** inset of the unrooted large subunit hydrogenase tree showing putative Group 3b hydrogenases in the CPR, along with presence/absence of HMM hits corresponding to other subunits **b)** genomic context for hydrogenase gene clusters, where position 0 corresponds to the location of the ORF encoding the large subunit of the NiFe hydrogenase. Only protein families on the same strand as the large subunit are represented in the charts, whereas genome diagrams below the charts include all proximal families regardless of strand orientation. **c)** Inset of large subunit tree within Group 4 hydrogenases. EHR: Energy-converting hydrogenases related complexes. Scale bars represent the average number of substitutions per site.

Supporting the possibility of transfer is the observation that multiple distinct enzyme forms underlie the distributions of several glycolytic functions. For example, we recovered unique hits to three individual HMMs representing various versions of PGI - one describing a general, cross-domain version (PF00342), another a bifunctional PGI/phosphomannose isomerase present in some bacteria and archaea (TIGR02128) (Hansen et al., 2004), and, finally, an unrelated cupin-based enzyme originally described from archaea (PF06560) (Hansen et al., 2005; Verhees et al., 2001). Interestingly, all three enzymes were scattered across the broad CPR groups, though very few CPR (∼2% of genomes) encoded more than one version. About 15 CPR genomes, mostly belonging to the Nealsonbacteria, encode only the cupin-related version. These sequences form a sibling clade to those from archaeal reference genomes in the corresponding gene tree (Supp. Fig. 4). CPR sequences with highest similarity to archaeal versions were also recovered for PGM (TIGR00306) and for TIM, although in the latter case sequences did not correspond to a separate HMM (Supp. Fig. 4). Similarly, while most CPR encode a class II FBA enzyme, some, particularly genomes belonging to the Kaiserbacteria and Woesebacteria, also encode a class I enzyme distinguished by a separate reaction mechanism (Cooper et al., 1996). Interestingly, in gene tree reconstructions for the class II aldolase, CPR sequences do not appear to be monophyletic, with small subgroups dispersed among sequences from bacterial reference genomes. Taken together, these results indicate that enzymes of multiple evolutionary origins underlie the distributions of core carbon metabolism, and support the idea that distributions have been shaped by episodes of lateral gene transfer, potentially from non-CPR bacteria or archaea.

### CPR bacteria encode phylogenetically distinct forms of Ni-Fe hydrogenases with variable genomic context

We next investigated the impact of lateral transfer on metabolisms sparsely distributed in the CPR, focusing on the NiFe hydrogenases that may play an important role in hydrogen economy and/or electron flux in this group as a case study (Castelle et al., 2018; Wrighton et al., 2012). Most CPR sequences were previously reported to fall within the Group 3b hydrogenases, cytoplasmic enzymes that may catalyze the reversible oxidation of H_2_ coupled to regeneration of NADPH or reduction of polysulfide when available (Silva et al., 2000; van Haaster et al., 2008). Here, a revised gene tree incorporating broadly sampled CPR reveals the presence of two subclades, which we term *hyd1* and *hyd2*, within CPR Group 3 hydrogenase (Fig. 4a). Both groups are related to, but distinct from, Group 3b versions in other bacteria and archaea, particularly *hyd2*, which is separated from its sibling clades by a relatively long branch (Fig. 4a).

Biochemically characterized Group 3b NiFe hydrogenases are known to be tetrameric enzymes (Pedroni et al., 1995). To examine whether subunit associations were consistent across CPR hydrogenase classes, we probed genomic context of the the large subunits using a paired HMM-protein clustering approach (Materials and Methods). Intriguingly, while both enzyme types generally encoded the small subunit hydrogenase (fam019) in addition to the catalytic subunit, only *hyd1* co-located with genes encoding protein families resembling the two other subunits involved in NAD(P)^+^-binding (gamma, fam034) and electron transfer (beta, fam012) (Fig. 4a). Consistent with previous work, HMM searches revealed that these subunits also have homology to anaerobic sulfide reductase A and B, suggesting that the entire complex could be involved in sulfur metabolism through the reduction of reduced sulfur compounds like polysulfide (Ma et al., 1993; Pedroni et al., 1995). However, in some cases, the gamma and beta subunits were not immediately upstream from the gene encoding the small subunit (Fig. 4b), and, in others, were not detected at all (Fig. 4a). This inconsistency may be potentially due to genome incompleteness or from lineage-specific losses within the *hyd1* clade.

Although genomes with *hyd2* also encoded the small subunit protein (fam019), the sequences were consistently truncated (mean 164 AA) relative to those associated with *hyd1* and non-CPR bacteria (mean 250 AA) (Supp. Fig. 5) (Vignais and Billoud, 2007). Both forms also shared fam002 in their genomic context, some members of which had homology to the hydrogenase-associated chaperone *hypC*. Outside these families, immediate genomic context differed for *hyd2*: while an HMM search recovered sequences with the NAD-binding motif (gamma subunit) in the vicinity of some *hyd2*, protein clustering showed that these proteins were neither proximal to, nor on the same strand as the catalytic subunit (Fig. 4ab). However, some members of fam013 that were in the genomic context of *hyd2* apparently possessed an NAD(P)-binding domain situated within a larger FAD-binding domain (PF07992). Similarly, while HMM searches did not recover evidence for a putative beta subunit near *hyd2*, we found one protein family (fam390) in proximity to a subset of *hyd2* that contained one of two iron-sulfur-binding domains. These domains were distinct from those associated with the putative beta subunit near *hyd1*. Ultimately, it is unclear whether *hyd2* consistently possesses (or lacks) the gamma and beta subunits and thus its function remains uncertain.

Intriguingly, both *hyd1* and *hyd2* were dispersed across many groups of CPR, and some CPR lineages contained both subtypes in closely related but distinct genomes (Fig. 4a). For example, genomes from the Roizmanbacteria, which harbored the largest total number of Group 3b-related NiFe hydrogenases (*n=15*), individually contained either *hyd1* or *hyd2* sequences. Mapping of genome taxonomy onto the 3b-related hydrogenase tree confirmed incongruencies with the CPR species tree (Fig. 4a). A similar pattern was observed for CPR sequences related to Group 4 NiFe hydrogenases that fell within a subclade of sequences representing energy-converting hydrogenases-related complexes (Ehr). Notably, these sequences were monophyletic and clustered separately from other Ehr proteins, although they also lacked the cysteine residues that bind the metal cofactors in other Group 4 enzymes (Supp. Fig. 5). This observation suggests that CPR Ehr proteins likely cannot interact with H_2_.

## DISCUSSION

Initially described as a radiation of phylum-level clades based on analyses of 16S rRNA divergence (Brown et al., 2015), the CPR has been thought to comprise at least 15% of bacterial phylum-level groups (Brown et al., 2015), potentially matching the scale of all other bacterial diversity (Hug et al., 2016). Attempts to adjust for lineage specific evolutionary rates have suggested the collapse of CPR into a single phylum (Parks et al., 2018), but more recent analyses with balanced taxonomic sampling continue to depict it as a substantial component of the bacterial domain (Zhu et al. 2019). Here, we combined new and previously reported genomes to construct a robust tree for the CPR radiation using two unlinked, concatenated marker sets (Fig. 1a, Supp. Fig. 3). The reconstructed trees are generally consistent with, and more clearly define, the topology originally described for the CPR (Brown et al., 2015), although definitive resolution of some deep nodes, particularly those connecting divergent groups like the Saccharibacteria, Gracilibacteria, and Absconditabacteria (SR1), remain elusive (possibly due to undersampling of the latter two lineages). Both gene trees support the presence of several monophyletic subgroups within the Parcubacteria, motivating subdivision of this large clade into smaller, taxonomically-relevant units.

Here, we evaluated metabolic platforms across the CPR by mapping genomically-encoded functions onto the concatenated ribosomal protein tree. Analysis of metabolic capacity among the CPR presents several challenges, primarily due to the fact that homologs of metabolic genes are often highly divergent compared to known reference sequences. Our custom approach for determining suitable cutoffs for HMMs indicate that manual threshold curation is important when proteins are only distantly related to biochemically characterized versions (Supp. Fig. 2,3). We found that metabolic platforms for CPR lineages only partially mirror phylogenetic relationships (Fig. 1ac), at least for the subset of metabolic traits examined here. In other words, phylogenetically distant lineages often possessed similar combinations of certain metabolic capacities (Fig. 1c). Thus, we hypothesize that diverse lineages of CPR may have converged upon similar metabolic platforms, potentially by specific combinations of lateral gene transfer and gene loss. This finding is intriguing, given that protein presence/absence patterns generally recapitulate phylogenetic relationships when entire proteomes are considered (Méheust et al., 2019). We thus infer that protein families not included in the current study must in balance show overall phylogenetic congruence to account for the observed difference.

Exploration of the patterns of gene distribution revealed that patchiness and phylogenetic depth varied for the selected metabolisms and even for enzymes in the same pathway (Fig. 2). As our analyses drew upon draft genomes (≥ 70% of genome markers present), it is possible that genome incompleteness impacted estimates of presence/absence for metabolic genes. However, given the large number and quality of genomes used in the analysis, we anticipate that these effects would only minimally alter the relative relationships between traits when examining depth and patchiness. While mis-binning can also complicate any analysis that relies upon metagenome-derived genomes, the similarity of findings for multiple closely-related genomes indicates that it likely does not greatly obscure the major patterns presented here. We used gene-species tree reconciliation to validate the prediction that proteins with variable ‘evolutionary profiles’ might have been shaped by different combinations of lateral transfer and vertical inheritance. For a subset of core carbon metabolism, here represented by glycolysis, gene trees were roughly congruent with the reconstructed organismal phylogeny, suggesting that vertical inheritance has primarily shaped distributions of these enzymes (Fig. 3). However, the discovery of a divergent subclade of TIM from CPR that is more closely related to archaeal versions than bacterial ones provides clear evidence of lateral transfer even for the most widely distributed glycolytic enzymes. Interestingly, two enzymes involved in the early steps of the glycolytic pathway (hexokinase/glucokinase and phosphofructokinase) were notably absent in nearly all lineages. Where present, they were likely acquired by lateral gene transfer, potentially following ancestral loss. These sequences separate from those of other bacteria, obscuring the source and suggesting that transfers of phosphofructokinase and hexokinase to CPR were also ancient. In contrast, enolase and pyruvate kinase, the last two steps of the pathway, are only somewhat widespread and show relatively low phylogenetic congruence. This pattern may reflect a mixture of genomic loss in addition to lateral transfer among unrelated CPR.

In archaea, glycolysis is known to be modified in a number of ways, including metabolic shunting (Imanaka et al., 2006) and rewiring of steps through novel enzymes (Siebers and Schönheit, 2005; Verhees et al., 2004). These observations have led to suggestions that evolutionary ‘tinkering’ has shaped glycolysis at least in some archaeal lineages (Van Der Oost and Siebers, 2007). Paralleling this, we found that several glycolytic steps in CPR are apparently carried out by different enzyme forms in different genomes, and, in some cases, by types that are traditionally associated with archaea. This was particularly striking in the case of PGI, which converts Glucose 6-P to Fructose 6-P, where three different enzyme forms accounted for the wide distribution of the function (Fig. 3a). Acquisition of variant enzymes may have preceded loss of the ancestral enzyme or occurred afterwards, complementing a loss in function. Overall, our findings suggest that CPR glycolysis is partly an evolutionary mosaic, as described in at least one eukaryotic organism (Stechmann et al., 2006), and further that gene loss and acquisition may have remodeled their glycolytic pathways over time.

Given the patchy distribution of enzymes involved in upper glycolysis, carbon flux through this portion of the pathway remains unclear. CPR without glucokinase/hexokinase (hex/glc) or PGI might rely on the uptake of glycolytic intermediates, like Fructose 6P or fructose 1,6-P2 from associated cells or released by cell lysis. These compounds could be shunted through the pentose phosphate pathway to bypass the largely absent phosphofructokinase and into the conserved central module of glycolysis (Fig. 3a) (Castelle et al., 2018). Alternatively, near universal conservation of TIM and GAPDH in the CPR suggests that either glycerone or G3P could also be important points of input for carbon flow in these organisms. Consistent with this is the fact that CPR encoding Form-III-related RuBisCO are predicted to introduce G3P to central/lower glycolysis as a product of their predicted nucleotide salvage pathway (Sato et al., 2007; Wrighton et al., 2016). The subset of CPR organisms that encode both hexokinase and PGI, on the other hand, could potentially perform a more diverse set of transformations, utilizing glucose precursors taken up from the environment or host. As for lower glycolysis, the observed patchiness in distributions of PGM, enolase, and pyruvate kinase suggests alternative fates for intermediates produced after the step catalyzed by PGK (Fig. 3a). In the absence of pyruvate kinase, which was found only in about a third of genomes here, CPR could use phosphoenolpyruvate (PEP) synthetase (PEPS) to instead interconvert PEP and pyruvate or instead generate oxaloacetate (Sauer and Eikmanns, 2005). Of course, with the data presented here, we cannot rule out the possibility that novel, divergent enzymes undetected by our HMM approach functionally substitute for those with patchy or nearly absent distributions among CPR. However, we found no evidence for the presence of archaeal PFK/glucokinase nor strong support for functioning of CPR ROK family proteins as putative glucokinases. Additionally, the CPR are not currently known to employ alternative pathways like the Entner-Doudoroff pathway, as do some other bacteria that lack PFK (Conway, 1992). Future work subjecting CPR in culture/co-culture to carbon flux analysis should help to validate genomic predictions and shed light on the metabolic configurations utilized *in vivo*.

Our second case study investigated the evolutionary history of specialized metabolism in the CPR bacteria, focusing on Group 4 and 3b NiFe hydrogenases (Fig. 4). These genes, like those putatively involved in nitrite reduction, electron transport, and AMP metabolism (Castelle et al., 2017; Danczak et al., 2017; Jaffe et al., 2019), are sparsely distributed among CPR and were likely subjected to lateral gene transfer. Notably, we report phylogenetic and genomic evidence for distinct monophyletic clades of Group 3b hydrogenases that are specific to the CPR. This suggests that transfer events were likely ancient or CPR hydrogenase sequences evolved very rapidly. The variable genomic contexts of the 3b-related *hyd1* and *hyd2* suggest at least two evolutionary scenarios: that individual, ancient transfers from non-CPR microorganisms occurred with the associated proteins intact, or that CPR encoding *hyd2* acquired only the large and small subunit and currently support function with unknown genes. The scattered distribution of both forms, phylogenetically incongruent with the CPR species tree, further suggests that intra-CPR exchange and/or loss also occurred over time. Similarly, we hypothesize that other sparsely distributed families in the CPR, like pyruvate:ferredoxin oxidoreductase, cytochrome oxidase, and nirK (nitrite metabolism), may also be the result of lateral transfer followed by further evolution within the CPR lineage. The acquisition of cytochrome oxidase by a small group of Saccharibacteria is presumably an adaptation to aerobic or microaerophilic environments (Castelle et al., 2018; Kantor et al., 2013; Starr et al., 2018).

In contemplating modes of evolution of CPR bacteria, it is important to consider the processes of gene gain and loss in the context of the largely symbiotic lifestyles of these organisms. The dynamic evolution of glycolysis might reflect reduced selection for complete pathways due to metabolic opportunities provided by the host, constraints which probably changed over time. Further, acquisition of new capacities via lateral transfer could have opened new niches, potentially including a change in or adaptation to new hosts in different environments. However, the observation that CPR sequences for rarer functions are often distinct from those of other bacteria suggests that these transfers probably occurred relatively early in the history of the radiation, or evolved rapidly once acquired. Distantly-related lineages within CPR may have independently undergone loss or gain of the same set of protein families, leading to similarly reduced metabolic platforms over time. These evolutionary constraints may be unique compared to those shaping minimal metabolism in other non-CPR bacterial groups with reduced genomes, like endosymbionts of insects. In contrast to these relatively recently evolved (linked to the appearance of eukaryotic hosts) associations that probably involve irreversible genome reduction trajectories (Moran and Wernegreen, 2000), the potential for CPR to associate with other bacteria raises the possibility of long-established symbioses in which gene sets remain in flux. The resulting pattern of ‘diversity within sparsity’ appears to be characteristic of the CPR.

## MATERIALS AND METHODS

### Genome collection and construction of phylogenetic marker sets

We compiled a large set of CPR genomes from metagenomes from several previous studies of various environments. We also binned an additional set of genomes from metagenomes previously generated from sediment from Rifle, Colorado (Anantharaman et al., 2016), groundwater from Crystal Geyser (Probst et al., 2017, 2018), a cyanobacterial mat from the Eel River network in northern California (Bouma-Gregson et al., 2019), and groundwater from a cold sulfide spring in Alum Rock, CA. Binning methods and taxonomic assignment followed Anantharaman et al. (2016). The total set was initially filtered for genomes that had been manually curated by any method to reduce the occurrence of misbinning, yielding a starting set of approximately 3,800 genomes. We next computed contamination and completeness for all genomes using a set of 43 marker genes sensitive to described lineage-specific losses in the CPR (Anantharaman et al., 2016; Brown et al., 2015) using the custom workflow in checkm (Parks et al., 2015). Results were then used to secondarily filter the genome set to those with >= 70% of the 43 marker genes present and <=10 % of marker genes duplicated. The resulting ∼2300 genomes were de-replicated at 95% ANI using dRep (*-sa 0*.*95 -comp 70 -con 10*) (Olm et al., 2017), yielding a set of 991 non-redundant genomes used for downstream analysis. These genomes along with their associated information, including accession numbers, are listed in Supp. Table 1.

We re-predicted for each genome using Prodigal (“single” mode) (Hyatt et al., 2010), adjusting the translation table (*-g 25*) for CPR lineages (Gracilibacteria and Absconditabacteria) known to utilize an alternative genetic code. Next, we assembled two sets of HMMs, representing the 16 syntenic ribosomal proteins (rp16) and, separately, the two subunits of RNA polymerase (RNAp), from the TIGRFAMs and Pfams databases and ran each against predicted proteins using HMMER v3.1b2 (http://hmmer.org) (Supp. Table 2). To maximize extracted phylogenetic information, including partial genes with robust homology to the marker genes, we set custom thresholds for each HMM using trees generated from all significant (e < 0.05) hits to a given HMM (aligned using MAFFT, tree inference with FastTreeMP) (Katoh and Standley, 2013; Price et al., 2010). Thresholds (listed in Supp. Table 2) were usually set at the highest bitscore attained by proteins outside the clade of interest (Supp. Fig. 1c), which were verified with BLASTp. HMM results and thresholds were visualized by in bitscore vs. e-value plots (Supp. Fig. 1ab). Phylogenetic analysis of HMM hits revealed that many proteins below model-specific thresholds were legitimate, often partial, hits to the targeted HMM (Supp. Fig. 1b).

Next, we curated phylogenetic marker sets for both rp16 and RNAp by addressing marker genes present in multiple copies in a given genomic bin. Multi-copy genes can result from remnant contamination after filtering, ambiguous bases in assembly leading to erroneous gene prediction (Parks et al., 2015), or legitimate biological features. We first identified marker genes fragmented by errors in gene prediction by searching for contiguous, above-threshold hits to the same HMM on the same assembled contig. This issue was particularly prevalent for rpoB and rpoB’, possibly due to repetitive regions in that gene impacting accurate assembly. For upstream fragments, we removed protein residues after stretches of ambiguous sequence to avoid introducing mis-translated bases into the alignment stage while maximizing phylogenetic information. If additional stretches of ambiguous sequence were present in downstream fragments, we removed them. Finally, we built a corrected, non-redundant marker set for each genome by selecting the 16 ribosomal proteins and, separately, 2 RNA polymerase subunits, that first maximized the number of marker genes on the same stretch of assembled DNA and, secondarily, maximized the combined length of encoded marker genes.

### Species tree inference, curation, and analysis

Results for each marker gene in the rp16 and RNAp sets were individually aligned with MAFFT (Katoh and Standley, 2013) and subsequently trimmed for phylogenetically informative regions using BMGE (-m BLOSUM30) (Criscuolo and Gribaldo, 2010). Gene trees for each marker were then constructed using IQTREE’s model selection and inference (*-m TEST -nt AUTO -st AA)* and manually inspected for major incongruencies.

In preparation for creating a concatenated alignment for each marker set, we next extracted corresponding rp16 and RNAp marker sets for a diverse bacterial outgroup consisting of ∼170 bacterial genomes from GenBank sampled evenly across characterized taxonomic divisions. We then merged the outgroup dataset with the existing CPR marker gene sets, individually aligning hits for each marker gene and trimming them as described above. We then concatenated individual protein alignments, retaining only those with both RNAp subunits and at least 8 of 16 syntenic ribosomal proteins. Maximum likelihood trees were inferred for both the concatenated rp16 (1427 AA) and RNAp (1652 AA) sets using ultrafast bootstrap and IQTREE’s extended FreeRate model selection (*-m MFP -st AA -bb 1500)* (Hoang et al., 2018; Kalyaanamoorthy et al., 2017; Nguyen et al., 2015), given the importance of allowing for site pattern heterogeneity in concatenated alignments (Wang et al., 2019). FASTA-formatted files for each marker, the unmasked alignment, the masked alignment, and newick trees for both rp16 and RNAp datasets are available in the Supplementary Material.

We next identified phylogenetic outliers in the resolved maximum likelihood topologies by searching for genomes that did not form a monophyletic clade with other organisms of the same taxonomy. These genomes, potentially due to mixed phylogenetic signal or undersampling, were retained only if they were assigned to a previously described novel lineage, or formed a conserved, uncharacterized clade with >1 member in both rp16 and RNAp trees. Genomes that did not fit these criteria were pruned. Concatenated trees were then re-inferred with the modified genome set. Where possible, we manually curated taxonomic assignment for genomes that clearly resolved within monophyletic clades of different taxonomic classification in both the rp16 and RNAp trees. Finally, we assessed broad-scale phylogenetic patterning within the CPR by examining the distribution of ribosomal proteins L1 and L9 employing the same HMM-based approach as described above.

### Metabolic annotation, analysis, and gene tree inference

To probe metabolism within the CPR, we assembled a broad set of HMMs from TIGRFAMs (tigrfams.jcvi.org/cgi-bin/Listing.cgi), Pfam (pfam.xfam.org), and a previous publication (Anantharaman et al., 2016) representing metabolisms relevant for biogeochemical cycling and energy production in this clade (Castelle et al., 2018; Kantor et al., 2013; Wrighton et al., 2012) (Supp. Table 2). We interrogated protein sequences from each CPR genome with the HMM set using HMMER and set custom bitscore thresholds as described above to ensure that divergent but functionally valid CPR proteins were retained. Model-specific thresholds were often much higher than maximum bitscores of CPR hits, even in cases where we were able to assign putative function to relatively high scoring clusters through BLAST and phylogenetic analyses. In a few cases (PRPP, PEP synthase, PGI, ROK family), we secondarily annotated HMM protein hits with additional Pfam domains or manually inspected placement within a reference tree to guide setting of accurate manual cutoffs. These additional domain HMMs and all custom thresholds are specific to this dataset and are listed in Supp. Table 2. If a protein had multiple above-threshold hits to a set of HMMs, we selected the HMM with the highest bitscore. We additionally selected the highest-scoring HMM hit within a genome bin for each HMM to generate a final set of metabolic markers for downstream analysis.

We next analyzed distributions of metabolic capacities in two ways: first, we created a presence/absence matrix for all metabolisms with at least one hit among the genome set, combining profiles for HMMs representing the same function (e.g. PGI, FBA, RuBisCO) into a single merged category. We then filtered the matrix to include only lineages with eight or more genomes and traits that were detected at least three times over all genomes. Finally, we averaged presence/absence across lineages, generating a frequency at which that trait was present among genomes of a particular taxonomy. We then used this information to generate a Bray-Curtis distance matrix using the ecopy package in Python. Finally, we performed a principal coordinates analysis using scikit-bio learn and plotted the resulting axes to examine clustering and variation within and among CPR metabolic platforms. Second, we measured phylogenetic conservation and patchiness over the rp16 tree using the consenTRAIT algorithm (*Npermutations = 1000, count_singletons = F, min_fraction = 0*.*90*) (Martiny et al., 2013) as implemented in the R package *castor* and consistency index (CI) as implemented in the R package *phangorn* and proposed in (*sitewise = T*) (Mendler et al., 2019). We integrated these two metrics to generate an “evolutionary profile” for each gene.

To build reference protein sets for the metabolic genes of interest, we queried proteins from the set of ∼170 bacterial reference genomes with same HMMs described above and applied the model-specific noise cutoff (for Pfam or TIGRFAMs HMMs) or the published cutoff (for custom HMMs). These proteins were then concatenated with the corresponding above-threshold hits from the CPR genomes and aligned as described above with MAFFT. Additionally, for four HMMs corresponding to glycolytic functions (PF06560, TIGR02128, TIGR00306, TIGR00419) we also queried a set of proteins from ∼300 archaeal reference genomes assembled in a similar fashion to the bacterial reference set. Resulting protein hits were concatenated with the bacterial sequences. For all single-gene alignments, columns with 95% or more gaps were trimmed using Geneious. Maximum-likelihood gene trees were then inferred using IQ-TREE with the following parameters: *-m TEST -st AA -bb 1500*. Trees were rooted on the largest monophyletic group of reference sequences present in the topology; if multiple monophyletic groups of reference sequences were present, trees were rooted at the midpoint.

To generate a gene tree for the NiFe hydrogenases, we assembled a comprehensive reference set of large subunit sequences from several published sources (Constant et al., 2011; Greening et al., 2016; Matheus Carnevali et al., 2019), dereplicated them at 95% amino acid identity using *usearch --cluster_fast*, and concatenated the resulting centroids with large subunit sequences recovered from the CPR. Sequences were aligned, alignments were trimmed, and the gene tree was inferred as described above for other metabolic genes. The sequence set, alignment, and trimmed alignment are available in the Supplementary Material. We next manually identified sequences within the immediate genomic context of 3b-related catalytic subunits that also scored highly against HMMs for anaerobic sulfite reductase A/B, as described previously for subunits in the Group 3b hydrogenases of *Pyrococcus furiosus* (Ma et al., 1993; Pedroni et al., 1995) and searched them for conserved domains in phmmer (https://www.ebi.ac.uk/Tools/hmmer/search/phmmer). We identified one iron-sulfur cluster and one NAD binding domain that were conserved among these proximal proteins (Supp. Table 2), and then queried all CPR proteins with these HMMs to identify putative 3b-related subunits across the entire genome set. We performed the same search for an additional Pfam domain associated with the 3b-hydrogenase small subunit (Supp. Table 2). For all three HMMs, manual thresholds were set using the paired visualization-phylogenetic approach described above. Finally, presence/absence of putative subunits were mapped onto the resolved tree of large-subunit sequences to examine patterns of association with phylogenetic clades of 3b-related hydrogenase using iTol (Letunic and Bork, 2016).

For the genomic context analysis of 3b-related forms, we gathered protein sequences within a 20 ORF radius (or less, if the scaffold ended) in both directions of the identified large subunits. Each ORF was assigned a genomic position relative to the large subunit (position 0). All recovered proteins were concatenated into a single file and passed through a two-part, *de novo* protein clustering pipeline recently applied to entire gene complements of CPR bacteria, in which proteins are first clustered into ‘subfamilies’ and highly similar/overlapping subfamilies are merged using and HMM-HMM comparison approach (--coverage 0.50) (Méheust et al., 2019) (https://github.com/raphael-upmc/proteinClusteringPipeline). Recovered protein families were compared with subunit HMM results and linked if the majority of proteins within the family had above-threshold hits to a given HMM. An alignment and gene tree for those proteins labelled as the small subunit hydrogenase (fam019) were made as described above.

Finally, counts for genes encoding the recovered families were plotted as a function of their relative position to the focal catalytic subunit of the hydrogenase across all CPR genomes. This was performed only if there were instances of the genes on the same strand (as predicted by Prodigal) as the large subunit hydrogenase. The relative positions of genes were multiplied by their strand orientation such that a negative position would signify being “upstream” of the focal catalytic subunit, whereas a positive position would signify being “downstream.” Positions were also adjusted in several cases were the focal subunit was split into multiple consecutive fragments, possibly due to local assembly errors.

### Data and software availability

Newly binned genomes used in the non-redundant, final set are available at **TBA**. Intermediate data files, including sequence files, and custom code used for the described analyses are available in interactive Jupyter Notebook format at https://github.com/alexanderjaffe/cpr-phylo-metab.

## Supporting information

supplementary trees/alignments

## ACKNOWLEDGMENTS

We thank Panagiotis Adam, Karthik Anantharaman, Raphaël Méheust, Najwa Taib, Daniela Megrian, Adi Lavy, Jacob West-Roberts, and Alexa Nicolas for informatics support and helpful discussions. Funding was provided by the Chateaubriand Fellowship to A.L.J., and the France Berkeley Fund to J.F.B and S.G.

## AUTHOR CONTRIBUTIONS

A.L.J. conducted the phylogenetic analyses, A.L.J., C.J.C., and P.M.C. performed the metabolic analyses. A.L.J., J.F.B., S.G., and C.J.C. developed the project. All authors contributed to the writing of the manuscript.

## SUPPLEMENTARY FIGURES

**Supp. Fig. 1.**
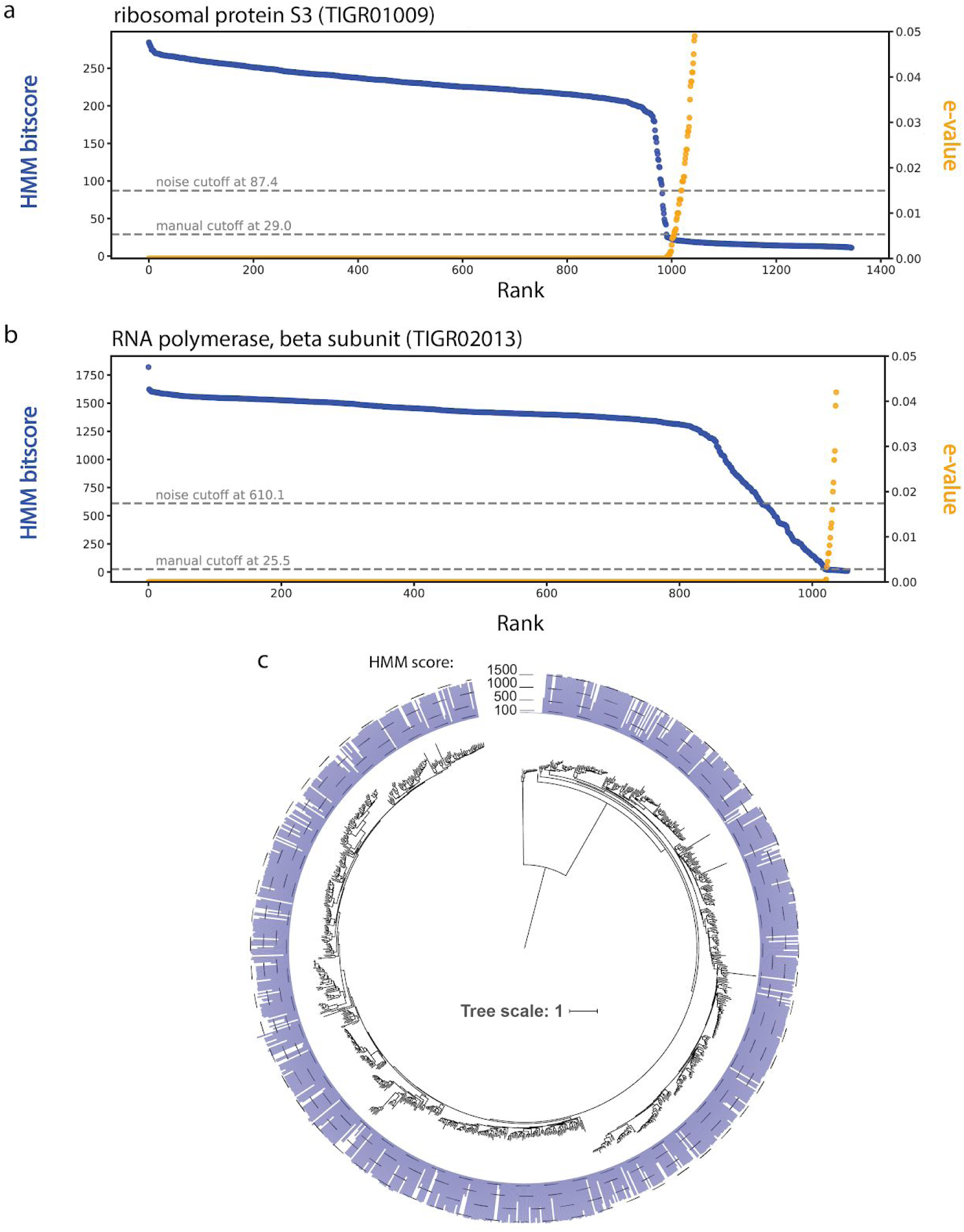
Visual and phylogenetic approach to setting sensitive manual thresholds for phylogenetic markers. HMM rank vs. bitscore/e-value plot for **a)** ribosomal protein S3 (TIGR01009) and **b)** RNA polymerase, subunit beta (TIGR02013). **c)** molecular phylogeny for significant (e > 0.05) TIGR02013 hits onto which HMM scores from **b)** are mapped.

**Supp. Fig. 2.**
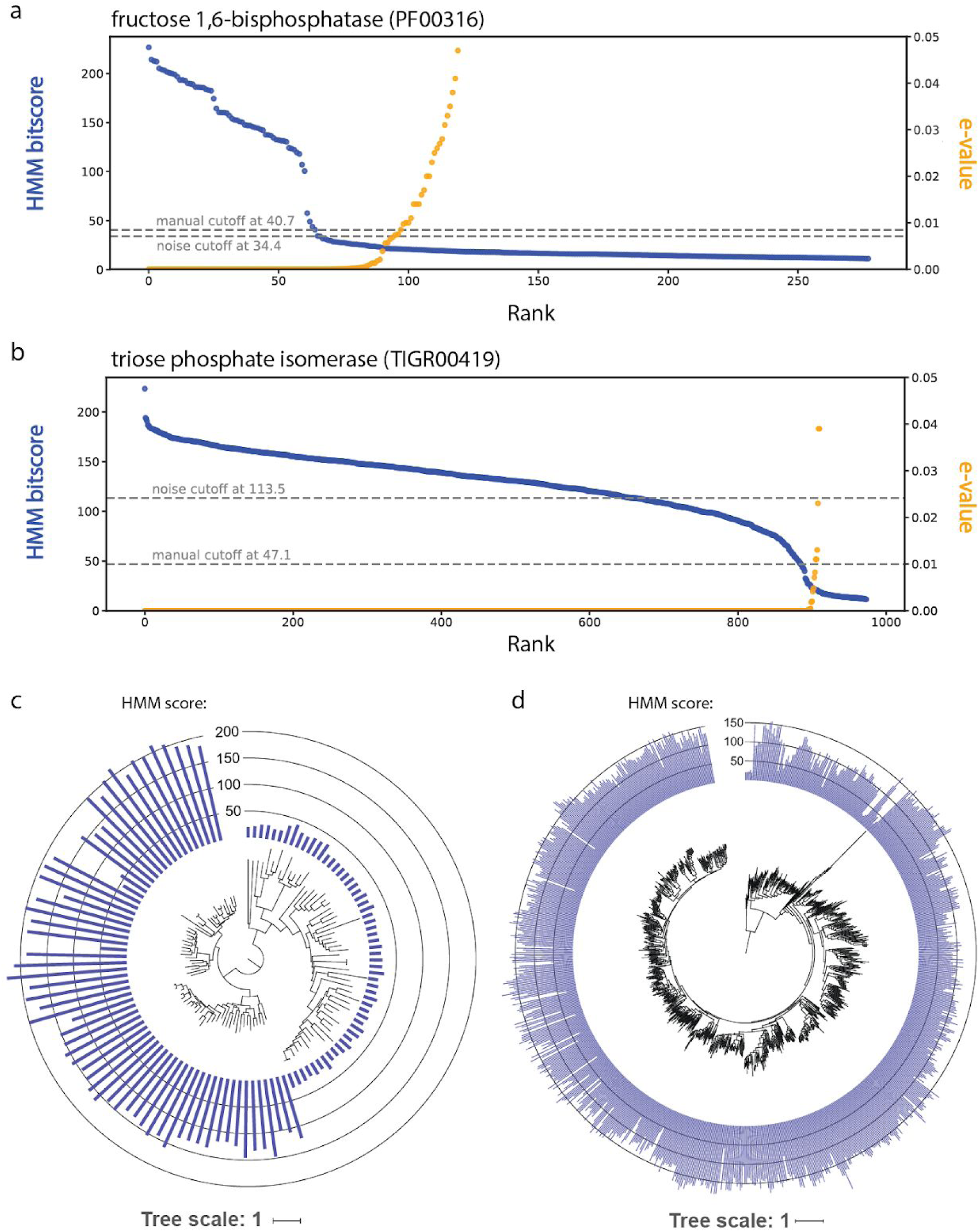
Visual and phylogenetic approach to setting sensitive manual thresholds for metabolic genes of interest. HMM rank vs. bitscore/e-value plot for **a)** fructose 1,6-bisphosphatase (PF00316) and **b)** triose phosphate isomerase (TIGR00419). Molecular phylogeny for significant (e > 0.05) hits to **c)** PF00316 and **d)** TIGR00419 onto which HMM scores are mapped.

**Supp. Fig. 3.**
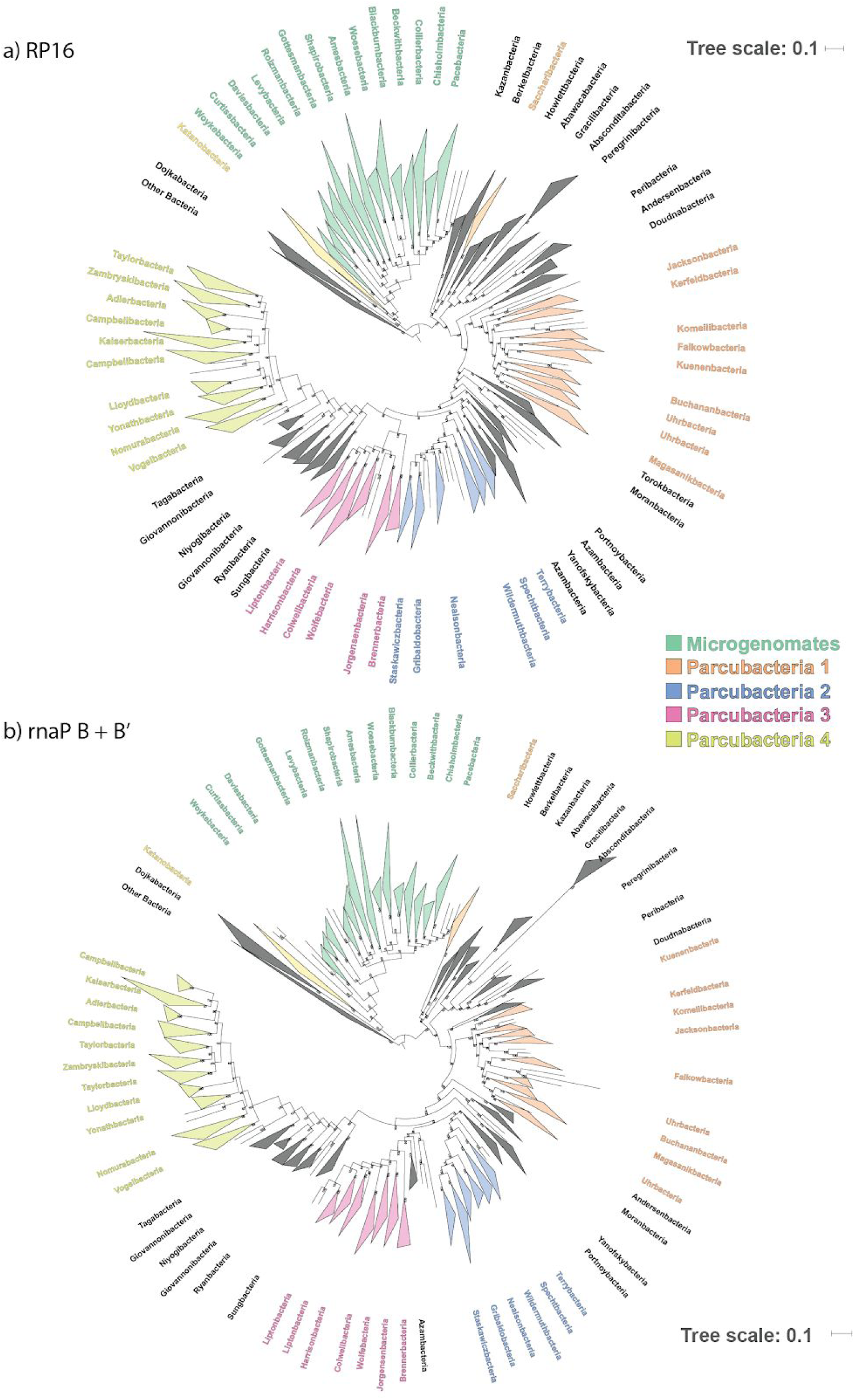
Consistent tree topology for CPR recovered individually by a concatenation of **a)** 16 ribosomal proteins and **b)** B and B’ subunits of RNA polymerase. Clade shading corresponds to that in Fig 1a. Scale bars represent the average number of substitutions per site. Ultrafast bootstrap support is indicated by the number attached to each tree node.

**Supp. Fig. 4.**
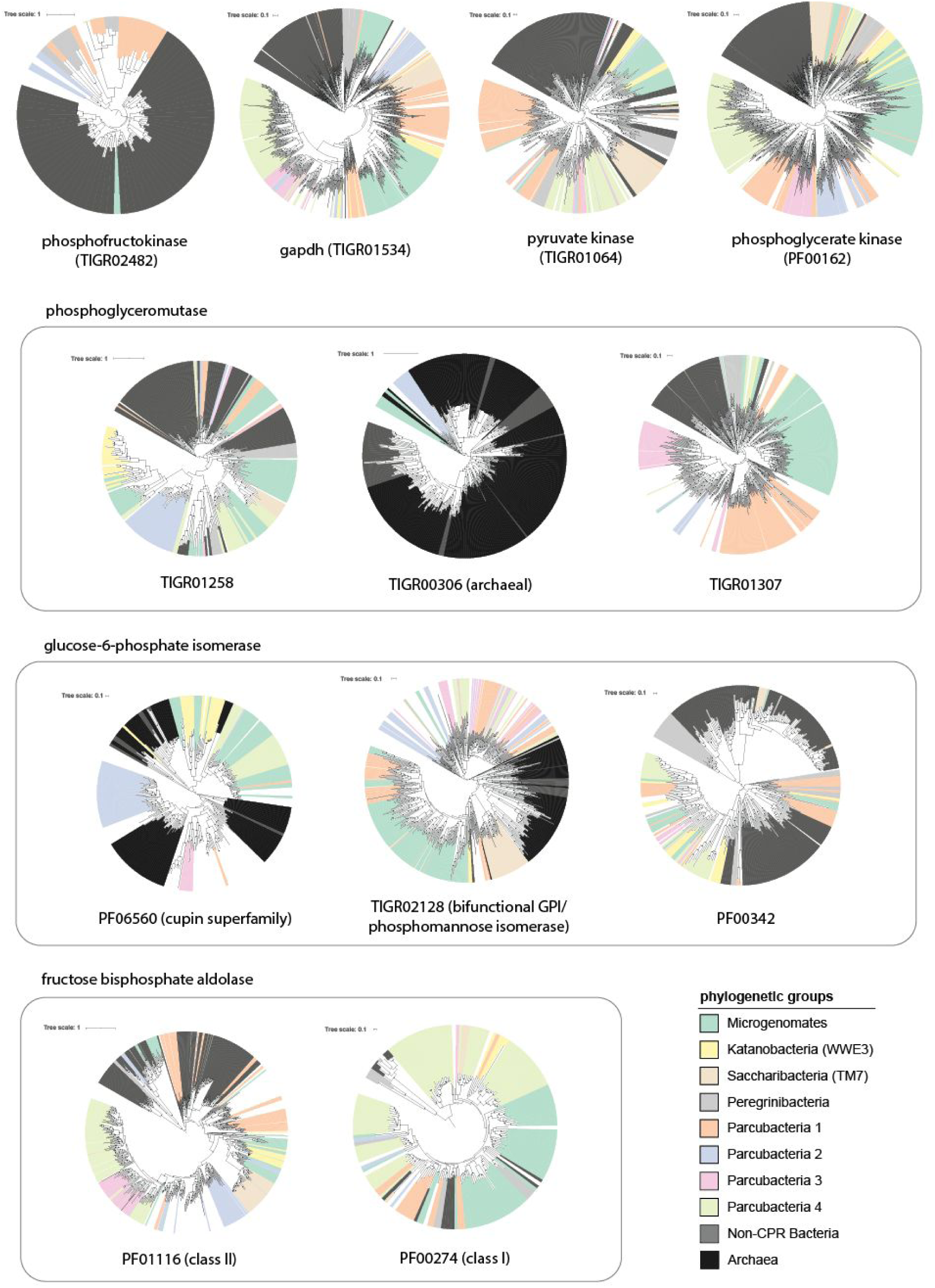
Maximum-likelihood gene trees for glycolytic enzymes in the CPR. Different HMMs representing the same functions are grouped together by boxes. Scale bars represent the average number of substitutions per site. Black dots indicate tree nodes with >=95% ultrafast bootstrap support.

**Supp. Fig. 5:**
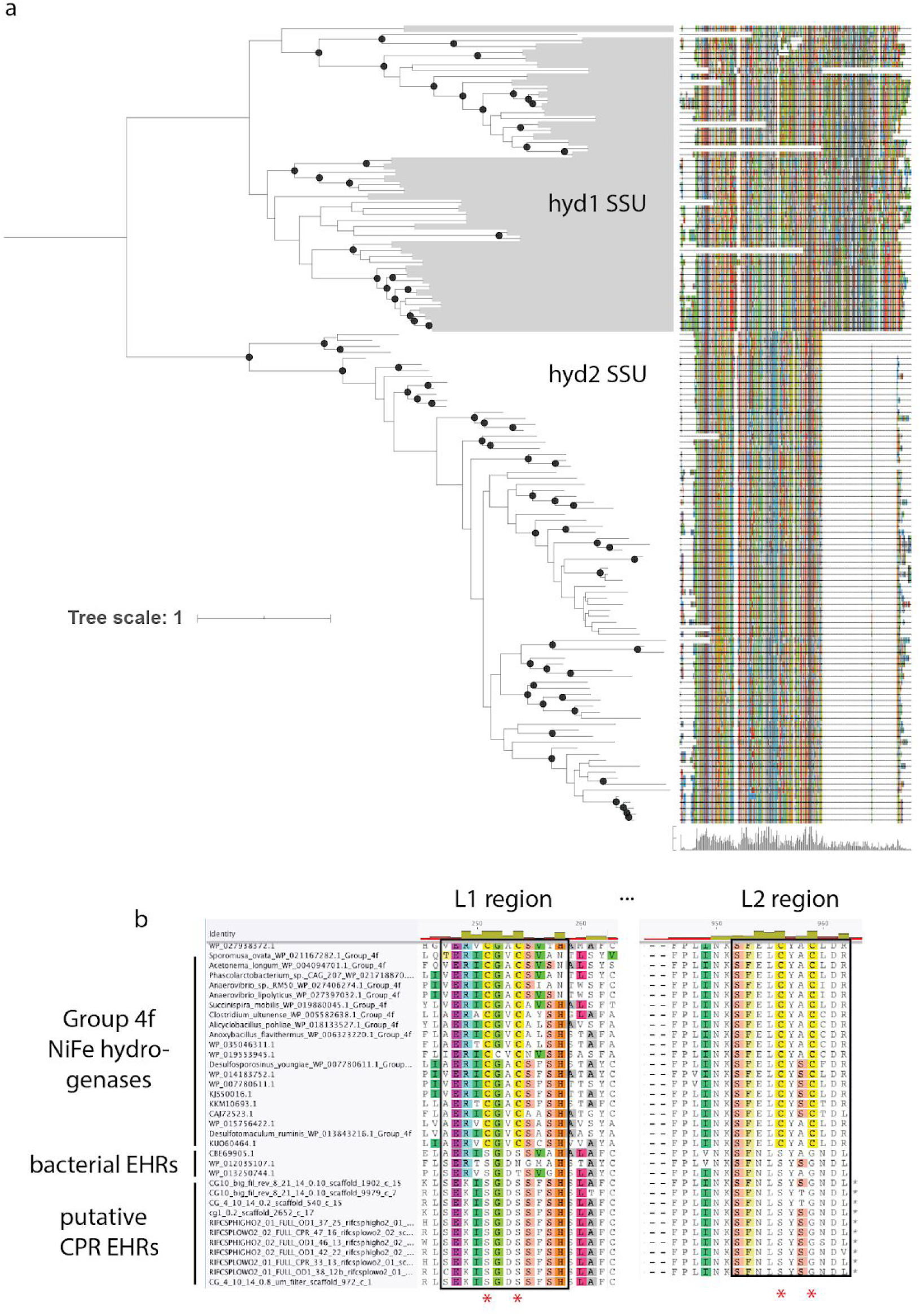
**a)** Maximum-likelihood gene tree for 3b-related NiFe hydrogenase small subunit (SSU) (fam019) with trimmed protein alignment for SSU sequences. Scale bar represents the average number of substitutions per site. Black dots indicate tree nodes with >=95% ultrafast bootstrap support. **b)** Partial alignment of the L1 and L2 regions of putative Group 4-related NiFe hydrogenases. EHR = energy-converting hydrogenases-related complexes. Red asterisk indicates cysteine residues associated with metal cofactor binding. **N**.**B**. for visual clarity, only a subset of sequences and sites are shown.

## SUPPLEMENTARY TABLES/FILES

**Supp. Table 1:** Characteristics of genomes used in this study.

**Supp Table 2:** Description of metabolic HMMs and thresholds used in this study.

**Supp. Material:** Trimmed alignments and inferred maximum-likelihood trees for the concatenated rp16 set, concatenated RNAp set, and NiFe hydrogenase large subunit.

## REFERENCES

Adam PS, Borrel G, Brochier-Armanet C, et al. (2017) The growing tree of Archaea: new perspectives on their diversity, evolution and ecology. The ISME journal 11(11): 2407–2425.

Anantharaman K, Brown CT, Hug LA, et al. (2016) Thousands of microbial genomes shed light on interconnected biogeochemical processes in an aquifer system. Nature communications 7: 13219.

Bouma-Gregson K, Olm MR, Probst AJ, et al. (2019) Impacts of microbial assemblage and environmental conditions on the distribution of anatoxin-a producing cyanobacteria within a river network. The ISME journal 13(6): 1618–1634.

Bräsen C, Esser D, Rauch B, et al. (2014) Carbohydrate metabolism in Archaea: current insights into unusual enzymes and pathways and their regulation. Microbiology and molecular biology reviews: MMBR 78(1): 89–175.

Brown CT, Hug LA, Thomas BC, et al. (2015) Unusual biology across a group comprising more than 15% of domain Bacteria. Nature 523(7559): 208–211.

Castelle CJ and Banfield JF (2018) Major New Microbial Groups Expand Diversity and Alter our Understanding of the Tree of Life. Cell 172(6): 1181–1197.

Castelle CJ, Brown CT, Thomas BC, et al. (2017) Unusual respiratory capacity and nitrogen metabolism in a Parcubacterium (OD1) of the Candidate Phyla Radiation. Scientific reports 7: 40101.

Castelle CJ, Brown CT, Anantharaman K, et al. (2018) Biosynthetic capacity, metabolic variety and unusual biology in the CPR and DPANN radiations. Nature reviews. Microbiology 16(10): 629–645.

Constant P, Chowdhury SP, Hesse L, et al. (2011) Genome data mining and soil survey for the novel group 5 [NiFe]-hydrogenase to explore the diversity and ecological importance of presumptive high-affinity H(2)-oxidizing bacteria. Applied and environmental microbiology 77(17): 6027–6035.

Conway T (1992) The Entner-Doudoroff pathway: history, physiology and molecular biology. FEMS microbiology reviews 9(1): 1–27.

Cooper SJ, Leonard GA, McSweeney SM, et al. (1996) The crystal structure of a class II fructose-1,6-bisphosphate aldolase shows a novel binuclear metal-binding active site embedded in a familiar fold. Structure 4(11): 1303–1315.

Criscuolo A and Gribaldo S (2010) BMGE (Block Mapping and Gathering with Entropy): a new software for selection of phylogenetic informative regions from multiple sequence alignments. BMC evolutionary biology 10: 210.

Danczak RE, Johnston MD, Kenah C, et al. (2017) Members of the Candidate Phyla Radiation are functionally differentiated by carbon- and nitrogen-cycling capabilities. Microbiome 5(1): 112.

Greening C, Biswas A, Carere CR, et al. (2016) Genomic and metagenomic surveys of hydrogenase distribution indicate H2 is a widely utilised energy source for microbial growth and survival. The ISME journal 10(3): 761–777.

Hansen T, Wendorff D and Schönheit P (2004) Bifunctional phosphoglucose/phosphomannose isomerases from the Archaea Aeropyrum pernix and Thermoplasma acidophilum constitute a novel enzyme family within the phosphoglucose isomerase superfamily. The Journal of biological chemistry 279(3): 2262–2272.

Hansen T, Schlichting B, Felgendreher M, et al. (2005) Cupin-type phosphoglucose isomerases (Cupin-PGIs) constitute a novel metal-dependent PGI family representing a convergent line of PGI evolution. Journal of bacteriology 187(5): 1621–1631.

Hoang DT, Chernomor O, von Haeseler A, et al. (2018) UFBoot2: Improving the Ultrafast Bootstrap Approximation. Molecular biology and evolution 35(2): 518–522.

Hug LA, Baker BJ, Anantharaman K, et al. (2016) A new view of the tree of life. Nature microbiology 1: 16048.

Hyatt D, Chen G-L, Locascio PF, et al. (2010) Prodigal: prokaryotic gene recognition and translation initiation site identification. BMC bioinformatics 11: 119.

Imanaka H, Yamatsu A, Fukui T, et al. (2006) Phosphoenolpyruvate synthase plays an essential role for glycolysis in the modified Embden-Meyerhof pathway in Thermococcus kodakarensis. Molecular microbiology 61(4): 898–909.

Jaffe AL, Corel E, Pathmanathan JS, et al. (2016) Bipartite graph analyses reveal interdomain LGT involving ultrasmall prokaryotes and their divergent, membrane-related proteins. Environmental microbiology 18(12): 5072–5081.

Jaffe AL, Castelle CJ, Dupont CL, et al. (2019) Lateral Gene Transfer Shapes the Distribution of RuBisCO among Candidate Phyla Radiation Bacteria and DPANN Archaea. Molecular biology and evolution 36(3): 435–446.

Kalyaanamoorthy S, Minh BQ, Wong TKF, et al. (2017) ModelFinder: fast model selection for accurate phylogenetic estimates. Nature methods 14(6): 587–589.

Kantor RS, Wrighton KC, Handley KM, et al. (2013) Small genomes and sparse metabolisms of sediment-associated bacteria from four candidate phyla. mBio 4(5): e00708–13.

Katoh K and Standley DM (2013) MAFFT multiple sequence alignment software version 7: improvements in performance and usability. Molecular biology and evolution 30(4). academic.oup.com: 772–780.

Letunic I and Bork P (2016) Interactive tree of life (iTOL) v3: an online tool for the display and annotation of phylogenetic and other trees. Nucleic acids research 44(W1): W242–5.

Luef B, Frischkorn KR, Wrighton KC, et al. (2015) Diverse uncultivated ultra-small bacterial cells in groundwater. Nature communications 6: 6372.

Ma K, Schicho RN, Kelly RM, et al. (1993) Hydrogenase of the hyperthermophile Pyrococcus furiosus is an elemental sulfur reductase or sulfhydrogenase: evidence for a sulfur-reducing hydrogenase ancestor. Proceedings of the National Academy of Sciences of the United States of America 90(11): 5341–5344.

Martiny AC, Treseder K and Pusch G (2013) Phylogenetic conservatism of functional traits in microorganisms. The ISME journal 7(4): 830–838.

Matheus Carnevali PB, Schulz F, Castelle CJ, et al. (2019) Hydrogen-based metabolism as an ancestral trait in lineages sibling to the Cyanobacteria. Nature communications 10(1): 463.

Méheust R, Burstein D, Castelle CJ, et al. (2019) The distinction of CPR bacteria from other bacteria based on protein family content. Nature communications 10(1): 4173.

Mendler K, Chen H, Parks DH, et al. (2019) AnnoTree: visualization and exploration of a functionally annotated microbial tree of life. Nucleic acids research 47(9): 4442–4448.

Moran NA and Wernegreen JJ (2000) Lifestyle evolution in symbiotic bacteria: insights from genomics. Trends in ecology & evolution 15(8): 321–326.

Nguyen L-T, Schmidt HA, von Haeseler A, et al. (2015) IQ-TREE: a fast and effective stochastic algorithm for estimating maximum-likelihood phylogenies. Molecular biology and evolution 32(1): 268–274.

Olm MR, Brown CT, Brooks B, et al. (2017) dRep: a tool for fast and accurate genomic comparisons that enables improved genome recovery from metagenomes through de-replication. The ISME journal 11(12). nature.com: 2864–2868.

Parks DH, Imelfort M, Skennerton CT, et al. (2015) CheckM: assessing the quality of microbial genomes recovered from isolates, single cells, and metagenomes. Genome research 25(7): 1043–1055.

Parks DH, Rinke C, Chuvochina M, et al. (2017) Recovery of nearly 8,000 metagenome-assembled genomes substantially expands the tree of life. Nature microbiology 2(11): 1533–1542.

Parks DH, Chuvochina M, Waite DW, et al. (2018) A standardized bacterial taxonomy based on genome phylogeny substantially revises the tree of life. Nature biotechnology. Nature Publishing Group, a division of Macmillan Publishers Limited. All Rights Reserved. DOI: 10.1038/nbt.4229.

Pedroni P, Della Volpe A, Galli G, et al. (1995) Characterization of the locus encoding the [Ni-Fe] sulfhydrogenase from the archaeon Pyrococcus furiosus: evidence for a relationship to bacterial sulfite reductases. Microbiology 141 (Pt 2): 449–458.

Price MN, Dehal PS and Arkin AP (2010) FastTree 2--approximately maximum-likelihood trees for large alignments. PloS one 5(3): e9490.

Probst AJ, Castelle CJ, Singh A, et al. (2017) Genomic resolution of a cold subsurface aquifer community provides metabolic insights for novel microbes adapted to high CO2 concentrations. Environmental microbiology 19(2): 459–474.

Probst AJ, Ladd B, Jarett JK, et al. (2018) Differential depth distribution of microbial function and putative symbionts through sediment-hosted aquifers in the deep terrestrial subsurface. Nature microbiology 3(3): 328–336.

Rinke C, Schwientek P, Sczyrba A, et al. (2013) Insights into the phylogeny and coding potential of microbial dark matter. Nature 499(7459): 431–437.

Sato T, Atomi H and Imanaka T (2007) Archaeal type III RuBisCOs function in a pathway for AMP metabolism. Science 315(5814): 1003–1006.

Sauer U and Eikmanns BJ (2005) The PEP-pyruvate-oxaloacetate node as the switch point for carbon flux distribution in bacteria. FEMS microbiology reviews 29(4): 765–794.

Schönheit P, Buckel W and Martin WF (2016) On the Origin of Heterotrophy. Trends in microbiology 24(1): 12–25.

Sieber CMK, Paul BG, Castelle CJ, et al. (2019) Unusual metabolism and hypervariation in the genome of a Gracilibacteria (BD1-5) from an oil degrading community. bioRxiv. DOI: 10.1101/595074.

Siebers B and Schönheit P (2005) Unusual pathways and enzymes of central carbohydrate metabolism in Archaea. Current opinion in microbiology 8(6): 695–705.

Silva PJ, van den Ban EC, Wassink H, et al. (2000) Enzymes of hydrogen metabolism in Pyrococcus furiosus. European journal of biochemistry / FEBS 267(22): 6541–6551.

Spang A, Saw JH, Jørgensen SL, et al. (2015) Complex archaea that bridge the gap between prokaryotes and eukaryotes. Nature 521(7551): 173–179.

Starr EP, Shi S, Blazewicz SJ, et al. (2018) Stable isotope informed genome-resolved metagenomics reveals that Saccharibacteria utilize microbially-processed plant-derived carbon. Microbiome 6(1): 122.

Stechmann A, Baumgartner M, Silberman JD, et al. (2006) The glycolytic pathway of Trimastix pyriformis is an evolutionary mosaic. BMC evolutionary biology 6: 101.

Tuininga JE, Verhees CH, van der Oost J, et al. (1999) Molecular and biochemical characterization of the ADP-dependent phosphofructokinase from the hyperthermophilic archaeon Pyrococcus furiosus. The Journal of biological chemistry 274(30): 21023–21028.

Van Der Oost J and Siebers B (2007) The glycolytic pathways of Archaea: evolution by tinkering. Archaea: evolution, physiology and molecular biology 22. Wiley Online Library: 247–260.

van Haaster DJ, Silva PJ, Hagedoorn P-L, et al. (2008) Reinvestigation of the steady-state kinetics and physiological function of the soluble NiFe-hydrogenase I of Pyrococcus furiosus. Journal of bacteriology 190(5): 1584–1587.

Verhees CH, Huynen MA, Ward DE, et al. (2001) The Phosphoglucose Isomerase from the Hyperthermophilic ArchaeonPyrococcus furiosus Is a Unique Glycolytic Enzyme That Belongs to the Cupin Superfamily. The Journal of biological chemistry 276(44): 40926–40932.

Verhees CH, Kengen SWM, Tuininga JE, et al. (2004) The unique features of glycolytic pathways in Archaea. Biochemical Journal. DOI: 10.1042/bj3770819.

Vignais PM and Billoud B (2007) Occurrence, classification, and biological function of hydrogenases: an overview. Chemical reviews 107(10): 4206–4272.

Wang H-C, Susko E and Roger AJ (2019) The Relative Importance of Modeling Site Pattern Heterogeneity versus Partition-wise Heterotachy in Phylogenomic Inference. Systematic biology. DOI: 10.1093/sysbio/syz021.

Wrighton KC, Thomas BC, Sharon I, et al. (2012) Fermentation, hydrogen, and sulfur metabolism in multiple uncultivated bacterial phyla. Science 337(6102): 1661–1665.

Wrighton KC, Castelle CJ, Wilkins MJ, et al. (2014) Metabolic interdependencies between phylogenetically novel fermenters and respiratory organisms in an unconfined aquifer. The ISME journal 8(7): 1452–1463.

Wrighton KC, Castelle CJ, Varaljay VA, et al. (2016) RubisCO of a nucleoside pathway known from Archaea is found in diverse uncultivated phyla in bacteria. The ISME journal 10(11): 2702–2714.

Zhu Q, Mai U, Pfeiffer W, et al. (2019) Phylogenomics of 10,575 genomes reveals evolutionary proximity between domains Bacteria and Archaea. Nature communications 10(1): 5477.

